# Memory encoding of syntactic information involves domain-general attentional resources. Evidence from dual-task studies

**DOI:** 10.1101/285411

**Authors:** Evelien Heyselaar, Katrien Segaert

**Affiliations:** Neurobiology of Language Department, Max Planck Institute for Psycholinguistics, Nijmegen, The Netherlands; School of Psychology, University of Birmingham, Birmingham, United Kingdom

**Keywords:** dual-task, attentional resources, language, syntactic priming, MOT

## Abstract

We investigate the type of attention (domain-general or language-specific) used during syntactic processing. We focus on syntactic priming: In this task, participants listen to a sentence that describes a picture (prime sentence), followed by a picture the participants need to describe (target sentence). We measure the proportion of times participants use the syntactic structure they heard in the prime sentence to describe the current target sentence as a measure of syntactic processing. Participants simultaneously conducted a motion-object tracking (MOT) task, a task commonly used to tax domain-general attentional resources. We manipulated the number of objects the participant had to track; we thus measured participants’ ability to process syntax while their attention is not-, slightly-, or overly-taxed. Performance in the MOT task was significantly worse when conducted as a dual-task compared to as a single task. We observed an inverted U-shaped curve on priming magnitude when conducting the MOT task concurrently with prime sentences (i.e., memory encoding), but no effect when conducted with target sentences (i.e., memory retrieval). Our results illustrate how, during the encoding of syntactic information, domain-general attention differentially affects syntactic processing, whereas during the retrieval of syntactic information domain-general attention does not influence syntactic processing.

## Introduction

Although Wundt in 1900 suggested that language requires attention (Wundt, 1900), most studies investigating the relationship between language and attention have only taken place in the last 20 years. As eye movements and attention are tightly coupled (Deubel & Schneider, 1996), eye gaze shifts and fixations are commonly used in language research as a real-time indicator of where the participant is attending at any given time. For example, studies on spoken word planning have shown that speakers tend to gaze at words and pictures until the completion of phonological encoding (e.g., Korvorst, Roelofs, & Levelt, 2006; Meyer, Sleiderink, & Levelt, 1998) and the seminal paper by Altmann & Kamide (1999) showed that listeners fixate on pictures before they are named, suggesting that we predict upcoming words based on the preceding words. Although these studies have provided evidence that language does require attention, it is still an open question as to what *kind* of attention is used. In this study we investigate whether syntactic processing uses domain-general or language-specific attentional resources.

There are suggestions that there is not one single pool of attentional resources (Kahneman, 1973; Wickens, 1980). Instead, dual-task studies have suggested that at least the visual and auditory domains rely on different attentional resources (Wickens, 2002). For example, Alais and colleagues (2006) illustrated nearly no effect on visual discrimination performance when participants performed a concurrent auditory chord and pitch discrimination task; however, performance decreased when the dual tasks were presented in the same modality. For language, there is no clear consensus on which attentional resources are necessary. It is likely that different aspects of language make different demands on attentional resources since central aspects of language are modality independent, and hence could instead tap into a domain-general “central executive” pool of attentional resources.

A core process of language production and comprehension is the processing of syntax. Syntax refers to the rules that assign grammatical roles and build phrase structure. There is no consensus (yet) on the steps involved in processing the syntax of a comprehended word/phrase (Friederici, 2002 vs. Hagoort, 2003). However, both models are based on ERP evidence which have suggested that some aspects, but not all, occur without the use of attention. The automaticity of syntax is supported by the fact that some steps occur very early (100 – 200ms after word onset; e.g. word category assignment), which is too fast for conscious, non-automatic control. Other steps in syntactic analysis occur later (300 – 600ms after word onset; e.g., morphosyntactic assignment) which is a long enough time period to include steps such as allocation of attentional resources in addition to the syntactic processing steps. However, to our knowledge, there have been no studies looking explicitly at whether syntactic processing requires attention, and if so, what type of attention is used (language-specific or domain-general). In this study we aim to shed light on this question.

A common method to measure the processing of syntax is via a syntactic priming task (Bock, 1986). In this task, the participants are exposed to frequently and infrequently used grammatical structures (e.g., *the man kisses the woman* vs *the woman is kissed by the man*) and the probability of the participant re-using the infrequent syntactic structure in their own utterances is used as a measurement of syntactic processing. This task has been used to test multiple characteristics of syntactic processing, such as the memory system used (Ferreira, Bock, Wilson, & Cohen, 2008; Heyselaar, Segaert, Walvoort, Kessels, & Hagoort, 2017) or how syntax is learned during development (Kidd, 2012).

In the current study, we aim to determine whether syntactic processing uses domain-general or language-specific resources. We aim to test this question by using a dual-task paradigm. If the performances of two simultaneously performed tasks are impaired, it suggests that the processing resources of these two tasks overlap. Hence increasing attention to one task almost always impairs performance on a second task (Kinchla, 1992), if they tap into the same resources. Otherwise there is no effect on secondary task performance. Dual-tasks are also used to support the structural interference theory, which posits that the human information-processing system can only select one independent response at one time (also known as the central bottleneck theory; Pashler, 1994). Hence by introducing a short time period between the start of Task 1 and the start of Task 2, one can measure how long the central processing stages of Task 1 take. The central bottleneck and resource allocation theories are not necessarily mutually exclusive (Temprado, Zanone, Monno, & Laurent, 2001). In our study, therefore, we will not give the participants the option on which task they can process first. We will ensure in our methodology that both tasks occur simultaneously, to prevent the participants from completing processing stages of one task before beginning the second. Therefore, by having participants conduct a secondary task during the syntactic priming task, we can manipulate the availability of attentional resources and measure how that effects syntactic processing.

For our concurrently presented task we will use a motion-object tracking (MOT) task (Pylyshyn & Storm, 1988), which relies on domain-general attention. In this task, participants are presented with a set of identical balls. A subset of these balls are briefly highlighted to indicate to the participants that they need to track the locations of these balls during the next phase of the task. The identical balls then move randomly around the screen for a set period of time. When they stop, the participant either has to indicate the location of the balls they were instructed to track or one ball is highlighted and the participant has to indicate whether this ball is one of the set they had to track. This task therefore requires attention throughout the entirety of a single trial (Scholl, 2008). By manipulating the number of balls the participant has to track, one can control the amount of attentional resources available for other, concurrent tasks. The MOT task has hence been used as a tool with which to manipulate domain-general attention in dual-task experiments (Allen, Mcgeorge, Pearson, & Milne, 2004; Fougnie & Marois, 2006; Postle, D’Esposito, & Corkin, 2005). Therefore, if there is an effect of doing this task concurrently with the language task, it is an indication that both tap into the same resources, suggesting that syntactic processing requires domain-general resources.

The classic comprehension-production syntactic priming task presents participants with a sentence describing a picture (prime trials), followed by a picture participants have to describe themselves (target trials). Due to the nature of the syntactic priming task, the prime trials tests language comprehension as the participants are listening to the picture descriptions, whereas the target trials tests language production as the participants are describing the picture. Therefore, we will run two separate experiments, one in which the MOT task is presented concurrently with the prime trials (hereafter named ‘Encoding phase’ as the dual-task is performed when participants encode the syntactic information) and one when it is presented concurrently with the target trials (hereafter name ‘Retrieval phase’ as the dual-task is performed when the participants retrieve the syntactic information). This will make it clearer when determining how attentional resources are used, as it could be that encoding requires more resources than retrieval.

Although syntax is an essential aspect of language, we predict that syntactic processing does require attention, and particularly domain-general resources due to the modality-independent nature of grammar processing. This would be reflected in our results as a decrease in priming magnitude with increasing attentional load. This study addresses the following questions: 1) does syntactic processing use domain-general resources, 2) how does syntactic processing respond to decreased attentional resources, and 3) is the interaction between syntactic processing and MOT task performance different depending on whether the syntactic information is encoded or retrieved?

## Methods

### Subjects

70 native Dutch speakers gave written informed consent prior to the experiment and were monetarily compensated for their participation. The participants were divided such that 35 participants completed the Encoding phase (10 male, M_Age_: 22.03 years, SD_Age_: 2.86) and the other 35 completed the Retrieval phase (10 male, M_Age_: 20.80 years, SD_Age_: 2.45). This study was approved by the ethics commission of the Faculty of Social Sciences at Radboud University, Nijmegen (Ethics Approval # ECG2013-1308-120).

### Statistical power

Statistical power was calculated using simulated priming data produced by the sim.glmm package (Johnson, Barry, Ferguson, & Müller, 2015) in R (R Core Development Team, 2011). For our simulated data set we assumed 20 repetitions per condition and 35 subjects. We assumed a 10% increase in passive production following a passive prime compared to baseline condition, as is commonly seen in the literature (Heyselaar, Hagoort, & Segaert, 2015; Segaert, Menenti, Weber, & Hagoort, 2011). With a difference of 6% between low ball load (low taxing of attention) and high ball load (high taxing of attention), our simulated data has a power of 0.878 with a 95% confidence interval of 0.856 - 0.898.

### Materials

#### Syntactic Priming Task

The pictures used in this task have been used elsewhere (Segaert, Menenti, Weber, & Hagoort, 2011). Our stimulus pictures depicted 40 transitive events such as *kissing, helping* or *strangling* with the agent and patient of this action. Each event was depicted by a greyscale photo containing either one pair of adults or one pair of children. There was one male and one female actor in each picture and each event was depicted with each of the two actors serving as the agent. To prevent the forming of strategies, the position of the agent (left or right) was randomized. These pictures were used to elicit transitive sentences; for each picture speakers can either produce an active transitive sentence (e.g. *the woman kisses the man*) or a passive transitive sentence (e.g. *the man is kissed by the woman*).

Filler pictures were used to elicit intransitive sentences. These fillers depicted events such as *running, singing,* or *bowing* using one actor. The actor could be any of the actors used in the transitive stimulus pictures. These intransitive sentences could be used as fillers, but also as a baseline measurement of each participant’s grammatical preferences. The intransitive picture would be used in the prime, with a transitive picture in the target, to measure how the participants would describe such sentences without being primed (baseline trial).

Each experimental list contained 24 targets in each of the 6 transitive priming conditions (active and passive prime for each of the 3 loads) and 24 targets in the baseline condition. We therefore have 24 repetitions for each condition. Within each experimental list, this resulted in 144 transitive descriptions on target pictures, 144 transitive descriptions on prime pictures and 72 intransitive descriptions leading up to a target in the baseline condition. The intransitive sentences also served as filler sentences in an extra 72 sentences. In total there were thus 432 sentences in the experiment. Over the whole experimental list 66% of the items (288 out of the total of 432 sentences) elicited transitive sentences.

### Task and Design

In order to manipulate the number of attentional resources available, participants completed a standard syntactic priming task and a motion-object tracking task (MOT task) simultaneously. Figure 1 depicts the order of events. The task was presented on a desktop computer using Presentation software (script available upon request), the recordings were played over headphones. The syntactic priming task used active (*the man kisses the woman*) or passive (*the woman is kissed by the man*) sentences. To aid understanding, we will describe the designs of each task separately and then describe how we combined them.

**Figure 1.**
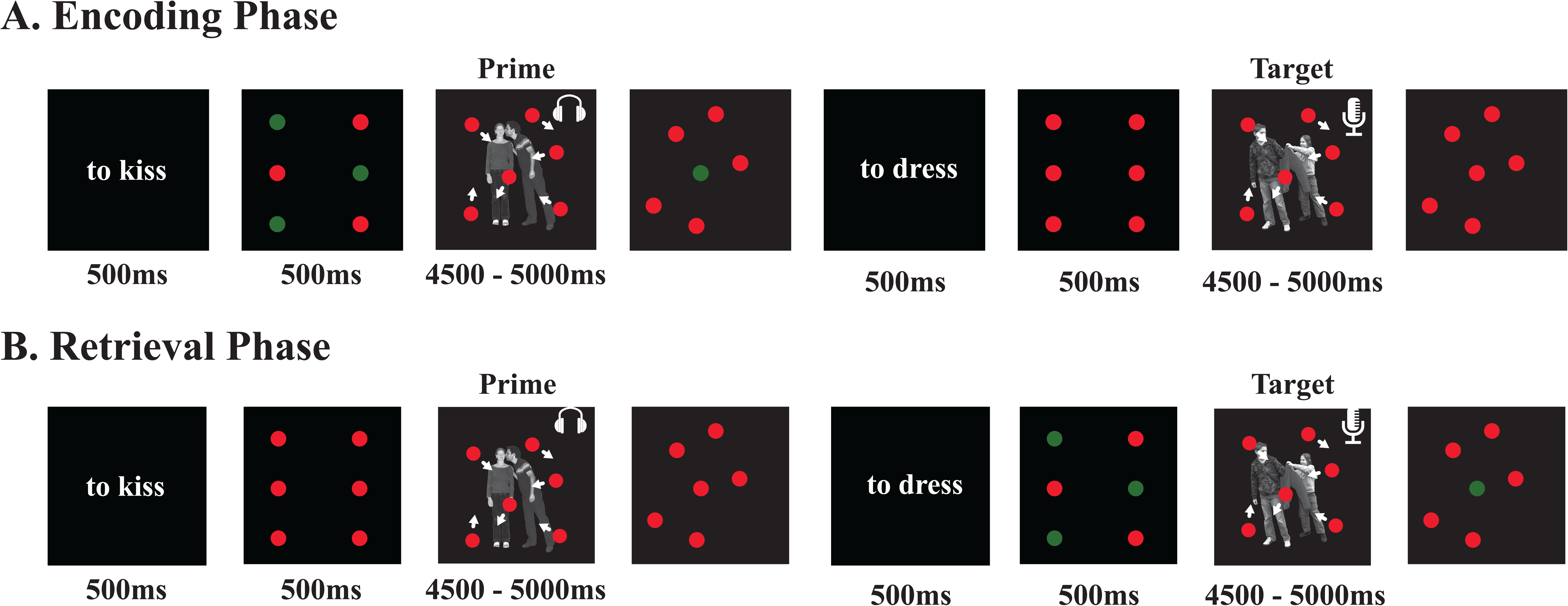
Experimental Design. Participants completed the dual task either in the Encoding phase (MOT task presented while participants listen to a picture description/prime phase of the priming task) or in the Retrieval phase (MOT task presented while participants describe a picture/target phase of the priming task). 0, 1 or 3 balls were briefly highlighted at the beginning of the MOT task that the participants have to track. Only one ball is highlighted at the end; participants respond via button press if this was one of the balls they had to track or not. If no balls are highlighted, the participants can effectively ignore the balls for the current trial. Ball load was randomized.

#### Syntactic Priming Task

Each trial consisted of a prime (participants listening to a recording) followed by a target (participants describing the picture using the verb provided). As mentioned above, a prime could be an active sentence (*the man kisses the woman*), a passive sentence (*the woman is kissed by the man*), or an intransitive/baseline sentence (*the man jumps*). A priming effect in our task is therefore defined as the proportion of passive sentences produced after hearing a passive prime, compared to the proportion of passive sentences produced after a baseline trial.

Participants were initially presented with a neutral verb (to be used in an upcoming utterance) for 500ms. After 500ms of black screen a greyscale picture would appear. Participants were instructed to either listen to a recording (presented 500ms after picture onset) which describes the picture, or describe the picture themselves using the neutral verb provided earlier. After 4500 – 5000ms (jittered) the picture is removed. The screen is black for an intertrial interval of 1500 - 2000ms (jittered) before the next verb is presented.

#### MOT Task

Participants are presented with a 2 by 3 array of 6 identically sized and shaped red balls. A subset of these (none, one or three) would be briefly highlighted green for 500 ms. After this they would all turn red again and start moving randomly around the screen. After 4500 – 5000 ms (jittered) the balls stop moving. One of the balls is highlighted green and the participant needs to indicate via key press whether that ball was one of the balls highlighted green at the beginning of the trial or not. If no balls were highlighted at the beginning, then no probe ball is highlighted at the end.

#### Dual Task

Each trial begins with the presentation of the neutral verb. During the 500ms wait time between verb presentation and picture presentation, the 2 by 3 array of balls would be presented, with the subset highlighted. The picture presentation and the ball movement initiation happened simultaneously to ensure no task started first. After 4500 – 5000ms (jittered) the balls stopped moving and the picture disappeared simultaneously. The intertrial interval of 1500 – 2000ms (jittered) will start once the participant has responded to the probe ball.

All participants completed prime and target trials. The participants who completed the dual-task in the Encoding phase would track balls during the prime only (so participants would track balls while listening to picture descriptions) whereas participants who did the dual-task in the Retrieval phase would track balls during the target only (participants would track balls while describing pictures). However, in both phases, both prime and target trials involved the presentation of moving balls to ensure the visual input is balanced between phases. No balls would be highlighted at the beginning of the trial, so participants knew they could effectively ignore the balls. The number of balls to track was randomized in one experimental session.

To ensure that participants paid attention to the recordings, 10% of the recordings did not match the picture on prime trials. The mismatch was balanced between role-switch of the agent and patient, incorrect verb used or incorrect agent/patient used. Participants were instructed to press a certain key if the recording was a mismatch.

### Procedure

Participants were informed that the experiment was about measuring multi-tasking ability. To ensure that the participants understood the task correctly, they first completed practice sessions of the MOT and syntactic priming task separately. The MOT practice session was used to calculate their baseline attentional capacity and contained 10 repetitions of each number of balls to track (0, 1 or 3). The syntactic priming task alone was too short to measure priming magnitude (at least 30 minutes is recommended for a stable effect, Heyselaar et al., 2015). No passives were used in the practice session to ensure participants were not primed before the main task began.

At the end of the practice session, participants could practice the MOT and syntactic priming task together to ensure they understood the order of events. This contained 5 prime-target trial pairs, of which none were passive structures. Participants could repeat this phase as many times as they wanted until they felt confident they knew how the tasks worked. No participant repeated the practice session more than thrice. Recent studies have shown that practice can reduce the psychological refractory period effect (Ruthruff, Johnston, Van Selst, Whitsell, & Remington, 2003; Van Selst, Ruthruff, & Johnston, 1999), again minimizing any central bottleneck influences in our study. During the actual experiment, the participant was given a short, self-timed break every 15 minutes to ensure motivation.

### Coding and analysis

Responses during the syntactic priming task were manually coded by the experimenter as either active or passive. Trials in which the descriptions did not match one of the coded structures were discarded. Target responses were included in the analysis only if 1) both actors and the verb were named (a sentence naming only one of the actors does not qualify as a transitive sentence) and 2) the structures used were active, passive or intransitive. In total 43 trials (0.57%) in the Encoding phase and 41 trials (0.55%) in the Retrieval phase were discarded.

The responses were analysed using a mixed-effect model, using the glmer and lmer functions of the lme4 package (version 1.1-4; Bates, Maechler, & Bolker, 2012) in R (R Core Development Team, 2011). Target responses were coded as 0 for actives and 1 for passives in the factor *Prime*. We used a maximal random-effects structure (Barr, Levy, Scheepers, & Tily, 2013; Jaeger, 2009): the repeated-measures nature of the data was modelled by including a per-participant and per-item random adjustment to the fixed intercept (“random intercept”). We attempted to include a maximal random effects structure; however, our model would not converge with all interactions (“random slopes”). We therefore reduced the structure by removing interactions before main effects. In terms of fixed effects, we began with a full model and then performed a step-wise “best-path” reduction procedure, removing interactions before main effects, to locate the simplest model that did not differ significantly from the full model in terms of variance explained. As we ran multiple models for each stage of analysis, details on each model are reported in the respective results section. For all models, the factor *Prime* was dummy coded so that we could determine how active and passive production compared to the baseline condition (reference group). *Phase* was sum contrasted and *Load* was reverse Helmert coded (current level compared to the mean of the preceding levels – a more accurate way to model linear relationship between levels). All numerical predictors were centred.

We also included the factor *Cumulative Passive Proportion*. This factor was calculated as the proportion of passives out of the total transitive responses produced on the target trials before the current target trial. A positive and significant *Cumulative Passive Proportion* therefore suggests that the proportion of passives previously produced positively influences the probability of producing a passive on the current target trial and is commonly used to model the learning effect of priming (Heyselaar et al., 2015; Jaeger & Snider, 2008; Segaert, Wheeldon, & Hagoort, 2016).

## Results

### Motion Object Tracking task

Figure 2 shows the behavioural results from the Motion Object Tracking (MOT) task. All participants completed the MOT task alone (without a secondary task). Half of the participants additionally completed the MOT task while listening to prime sentence descriptions (MOT+Encoding) and the other half of the participants completed the MOT task while describing the target picture (MOT+Retrieval). We observed no significant difference in the performance in the MOT Alone condition between the two participant groups (F(1,136) = 1.53, *p* = .219).

**Figure 2.**
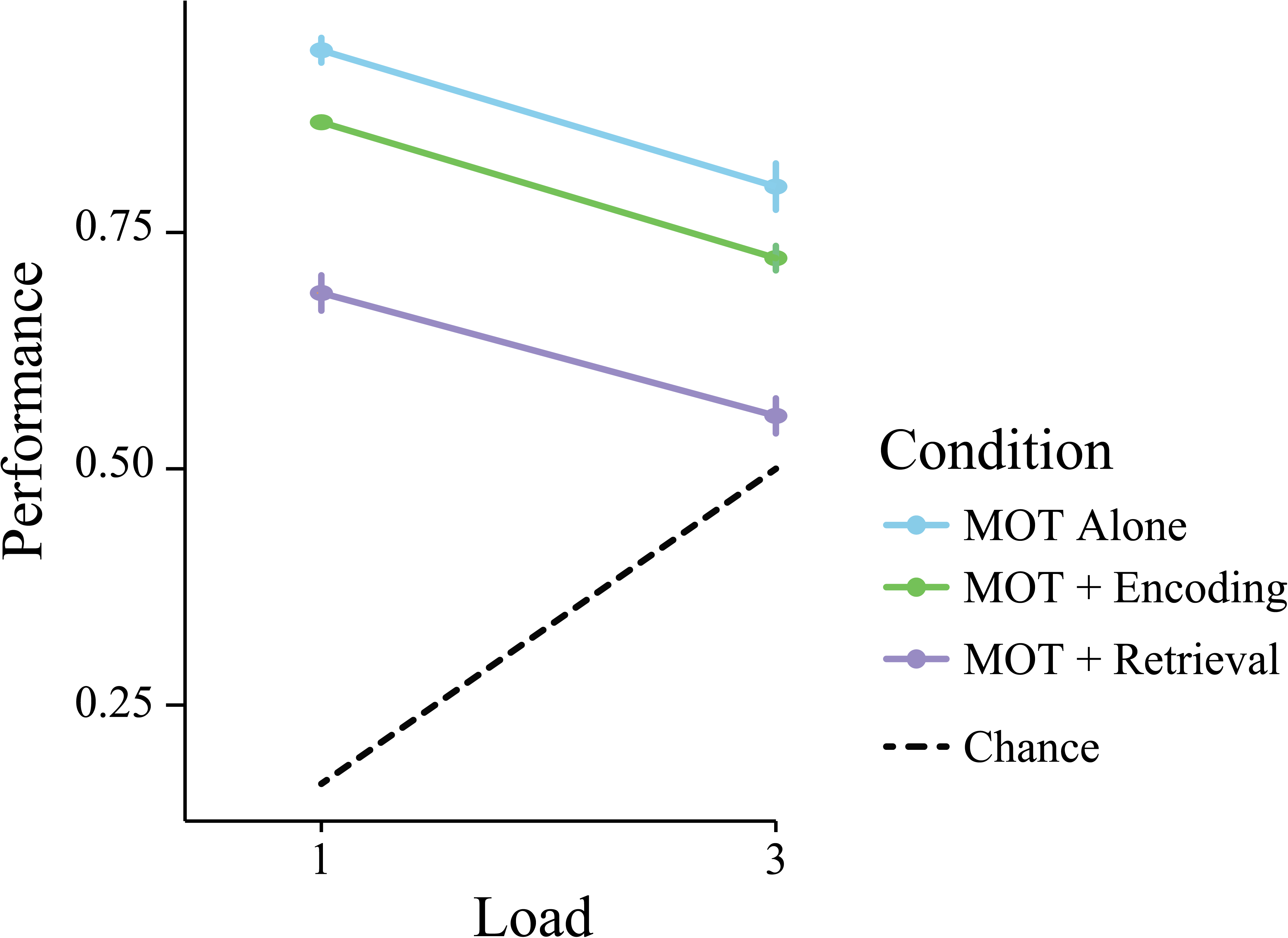
Motion Object Tracking Task (MOT) Performance. There is a significant difference in the proportion of correct responses between the different conditions compared to performing the MOT task alone. There was a greater drop in performance for the MOT + Retrieval condition than the MOT + Encoding condition. Error bars represent standard error.

A 3 (MOT Condition: Alone, +Encoding, +Retrieval) x 2 (MOT Load: 1 or 3 balls) between-subjects ANOVA revealed that MOT performance was reduced with increasing loads (F(1,274): 114.13, *p* < .001). More importantly, there was a main effect of condition (F(2,274): 135.85, *p* < .001) showing that performance on the MOT task was significantly reduced when conducted alone compared to in a dual-task scenario. This is consistent with previous dual-task literature: Performance of a single task is significantly better than performance of the same task in a dual-task scenario (Bourke, 1996).

Planned comparisons (Tukey’s HSD) revealed that, as illustrated in Figure 2, both the MOT+Encoding condition and the MOT+Retrieval condition were significantly different from the MOT Alone condition (*p* < .001). Interestingly, performance in the MOT+Retrieval condition was significantly worse compared to the MOT+Encoding condition (*p* < .001).

### Syntactic Priming Task

#### Single Task Effects

Firstly, we performed a logit mixed model on the Load (0) condition, as this is the equivalent of doing the syntactic priming task as a single task. We did this to ensure that our task did elicit a significant priming effect before we investigated whether the attentional manipulation influenced the magnitude of this effect. This model included *Prime*Phase* and *Phase*Cumulative Passive Proportion* as fixed effects, and as random slopes for the per subject intercept. The per item intercept took no random slopes.

We observed no significant difference in performance between the two phases in the Load (0) condition (β = 0.44, *p* = 0.147). As predicted, there is a significant influence of passive prime (β = 0.85, *p* = .011; 3.67% more passives following a passive prime compared to baseline – this is an average percentage for the Encoding and Retrieval Phase). This indicates that participants primed in our experiment. We additionally observed a significant influence of *Cumulative Passive Proportion* on passive target production (β = 9.51, *p* < .001). *Cumulative Passive Proportion* was calculated as the proportion of passives out of the total transitive responses produced on the target trials before the current target trial. A positive and significant *Cumulative Passive Proportion* therefore suggests that the proportion of passives previously produced positively influences the probability of producing a passive on the current target trial and is commonly used to represent the learning effect of priming (Heyselaar et al., 2015; Jaeger & Snider, 2008, 2013; Reitter, Keller, & Moore, 2011; Segaert et al., 2016). There was no effect of active primes (β = −0.65, *p* = .152, −1.48% on average between the Encoding and Retrieval Phase). We are therefore confident that our task elicits the same priming behaviour as seen in the literature in the absence of an attentional load manipulation (i.e. reverse preference effect and cumulativity; Jaeger & Snider, 2008, 2013; Reitter, Keller, & Moore, 2011; Segaert, Wheeldon, & Hagoort, 2016).

As our task elicited a robust priming effect akin to the magnitude seen in other studies, we are now able to investigate whether attentional load influenced the magnitude of this effect in the dual-task conditions.

#### Dual Task Effects

Both dual-task conditions (Encoding phase and Retrieval phase) contained prime-target pairs. During the prime the participant listened to a description of the picture, while during the target the participant described the picture. The only difference in conditions is that for the Encoding phase participants additionally had to complete the MOT task while listening to the prime picture; while for the participants in the Retrieval phase, they completed the MOT task while describing the target picture.

##### Catch Rate

To ensure that participants paid attention to the recordings, 10% of the recordings did not match the picture. These recordings were played only during the prime portion of the task and hence during the Encoding phase participants listened to the recordings while simultaneously completing the MOT task. During the Retrieval phase, the participants listened to the recordings in a single-task setting, as the MOT task was only presented in the target portion (when the participant describes the picture).

The catch rate was 95.2% (SD: 7.1%) and 91.1% (SD: 6.8%) for the Encoding and Retrieval phases respectively. Neither catch rate was significantly different from what is expected (χ^2^(2, N = 653) = 5.01, *p* = .082 for the Encoding phase, χ^2^(2, N = 630) = 2.18, *p* = .336 for the Retrieval phase). False alarm rate was 0.8% (SD: 0.48%) and 0.2% (SD: 0.00%) for the Encoding and Retrieval phases respectively. This indicates that even in a dual-task situation (Encoding phase), the participants still listened to the recordings to the same extent as in the single-task situation (Retrieval phase). The results are illustrated in Figure 3A.

**Figure 3.**
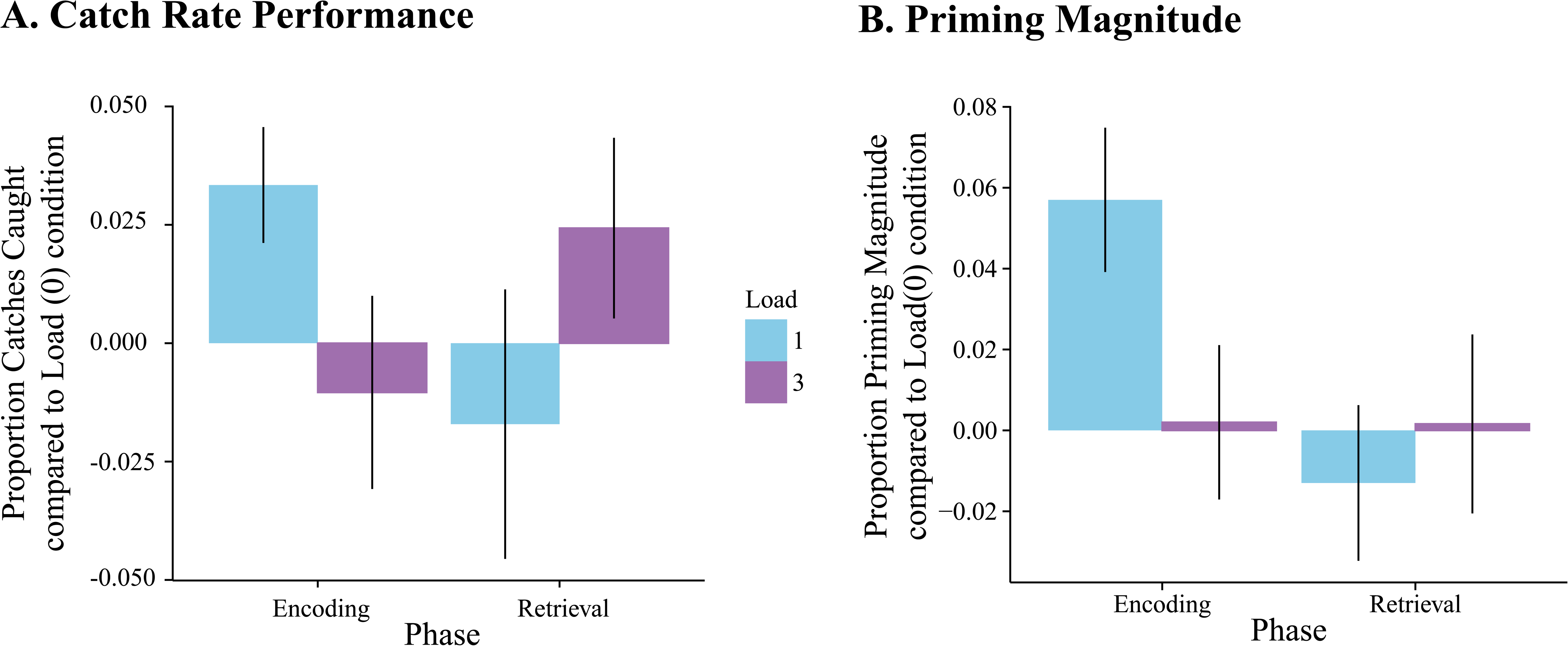
**A. Catch Rate per phase per load compared to Load (0) condition.** The figure illustrates the catch rate performance at each ball load compared to the no ball load (i.e. single-task) condition for each phase respectively. Note that there was no dual-task condition for any of the prime trials in the Retrieval phase. **B. Priming magnitude per phase per load compared to Load (0) condition.** The figure illustrates the amount of priming magnitude difference at each ball load compared to the no ball load (i.e. single-task) condition for each phase respectively. This better illustrates the effect the dual-task scenario has on performance. There is a significant difference in passive priming magnitude between phases (*p* = .026) as well as a Prime by Phase by Load interaction. Error bars represent standard error.

As the chi-squared test for the Encoding phase had a p-value of .082, we aimed to see whether the catch rate of the Load (1) condition was significantly higher compared to the Load (0) and Load (3) condition (as is suggested by Figure 3A). Indeed, this is the case (χ^2^(1, N = 653) = 4.70, *p* = .030).

##### Priming Effects

Figure 3B shows the priming magnitude for each ball load, for each phase, compared to the Load (0) condition. Priming magnitude was calculated as the proportion of passive responses after a passive prime compared to passive responses after an intransitive (not primed) sentence (baseline condition). As stated in *Single Task Effects*, the average priming magnitude for the Load (0) condition was 3.67% passive priming, a low yet robust effect (*p* = .011). We observed a 7.10% and 4.20% passive priming magnitude for Load (1) for the Encoding and Retrieval Phases, respectively, and a 1.60% and 3.90% passive priming magnitude for Load (3) for the Encoding and Retrieval Phases respectively. The figure illustrates the difference in priming magnitude at each ball load compared to the no ball (i.e., single-task) condition to better illustrate how the priming magnitude differed compared to the Load (0) condition.

The priming data was analysed using a logit mixed model. We began with a full model (*Prime*Load*Phase*Cumulative Passive Proportion*), and then performed a step-wise “best-path” reduction procedure, removing interactions before main effects, to locate the simplest model that did not differ significantly from the full model in terms of variance explained (Full = AIC: 4447.2, BIC: 4864.5; Best = AIC: 4435.1, BIC: 4728.0; *p* = .187). Multicollinearity was acceptable (VIF < 3.17), suggesting minimal Type II error from including factors that correlate to each other. The model took *Prime* and *Load* as random slopes for the per participants random intercept, and *Load* for the per item random intercept. The fixed effects of the best model fit for these data are summarized in Table 1.

**Table 1.** Summary of the fixed effects in the mixed logit model for the response choices based on prime structure.

| Predictor | coefficient | *SE* | *Wald Z* | *p* |  |
| --- | --- | --- | --- | --- | --- |
| Intercept | -3.97 | 0.24 | -16.70 | < .001 | *** |
| Active Prime | -0.37 | 0.23 | -1.65 | .099 |  |
| Passive Prime | 0.87 | 0.19 | 4.16 | < .001 | *** |
| Load (1) | -0.06 | 0.13 | -0.45 | .653 |  |
| Load (3) | 0.06 | 0.07 | 0.83 | .407 |  |
| Phase | 0.27 | 0.20 | 1.37 | .170 |  |
| Cum. Passive Proportion | 4.94 | 0.68 | 7.29 | < .001 | *** |
| Active Prime * Load (1) | -0.21 | 0.15 | -1.37 | .170 |  |
| Passive Prime * Load (1) | 0.07 | 0.12 | 0.57 | .570 |  |
| Active Prime * Load (3) | -0.02 | 0.08 | -0.30 | .764 |  |
| Passive Prime * Load (3) | -0.09 | 0.07 | -1.24 | .215 |  |
| Active Prime * Phase | 0.13 | 0.13 | 1.00 | .316 |  |
| Passive Prime * Phase | -0.25 | 0.13 | -1.90 | .058 |  |
| Phase * Load (1) | -0.23 | 0.10 | -2.25 | .024 | * |
| Phase * Load (3) | 0.02 | 0.05 | 0.44 | .660 |  |
| Active Prime * Phase * Load (1) | 0.33 | 0.15 | 2.20 | .028 | * |
| Passive Prime * Phase * Load (1) | 0.29 | 0.12 | 2.39 | .017 | * |
| Active Prime * Phase * Load (3) | -0.03 | 0.08 | -0.34 | .733 |  |
| Passive Prime * Phase * Load (3) | -0.03 | 0.07 | -0.40 | .691 |  |

Note: N = 11176, log-likelihood = -2177.6
\*\*\* < .001 \* < .05

The model shows a significant influence of passive primes on passive production (*p* < .001) and no significant influence of active primes on active production (*p* = .099). There is also a significant influence of *Cumulative Passive Proportion* on passive production. This again repeats the basic priming effects seen in the Load (0) condition reported above.

In terms of the aims of this study, there is a three-way interaction between Prime (Active or Passive), Phase (Encoding or Retrieval) and Load (0, 1 or 3 balls tracked). Passive Prime by Encoding Phase by Load (1) was significant (*p* = .017). To better understand the nature of this three-way interaction, we reanalysed the data per condition using logit mixed models. The results of these models are summarized in Table 2.

**Table 2.**
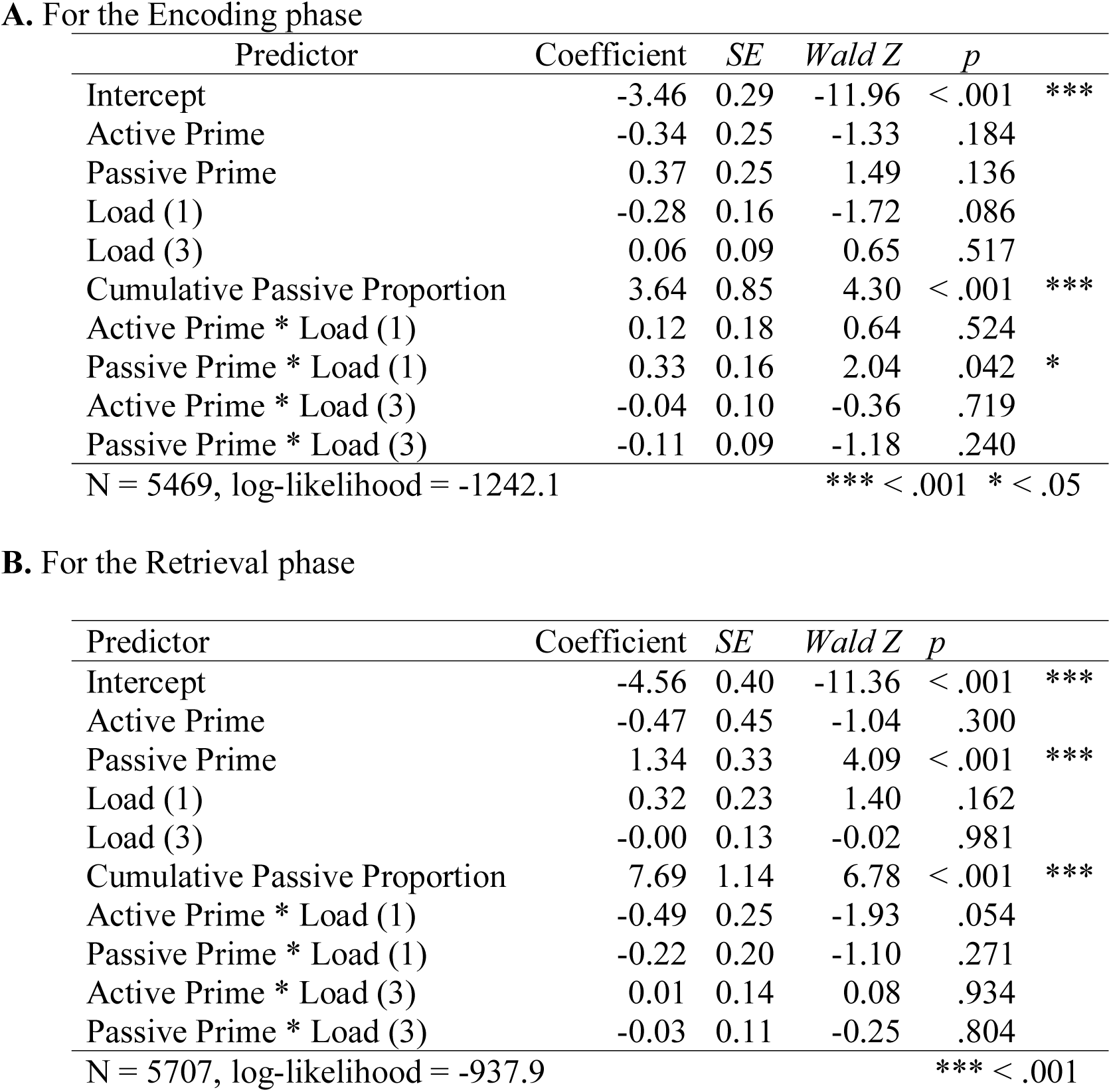
Summary of the fixed effects in the mixed logit model for the response choices based on prime structure and load.

Table 2A and B show that the three-way interaction from Table 1 is driven by a significant effect of holding one ball in attention during the Encoding phase on passive priming magnitude (*p* = .042). This effect is not seen for active priming (*p* = .524). We also see a trend towards there being slightly more active priming in the Load (1) condition for the Retrieval phase (*p* = .054).

This is similar to what was seen for the Catch Rate performance, suggesting that there is a boost in performance in both memory for the syntactic structure (as illustrated by the increase in priming magnitude) as well as the integration of audio and visual streams (as illustrated by the increased catch rate).

### Syntactic Priming and MOT

We correlated the task performance in the single MOT task condition with priming magnitude, to determine if being good at one task predicts individual performance in the other. A correlation would suggest that the relationship we have found between the tasks may not be due to shared attentional resources, but due to the fact that some individuals are better that attention-dependent processes, such as goal maintenance and/or persistence.

We show no correlation between the task performance for the Encoding phase (Spearman’s rho = 0.079, *p* = .521; Figure 4) nor for the Retrieval phase (Spearman’s rho = −0.093, *p* = 442). Therefore, we are confident that the interaction we see is truly because they tap into the same resources.

**Figure 4.**
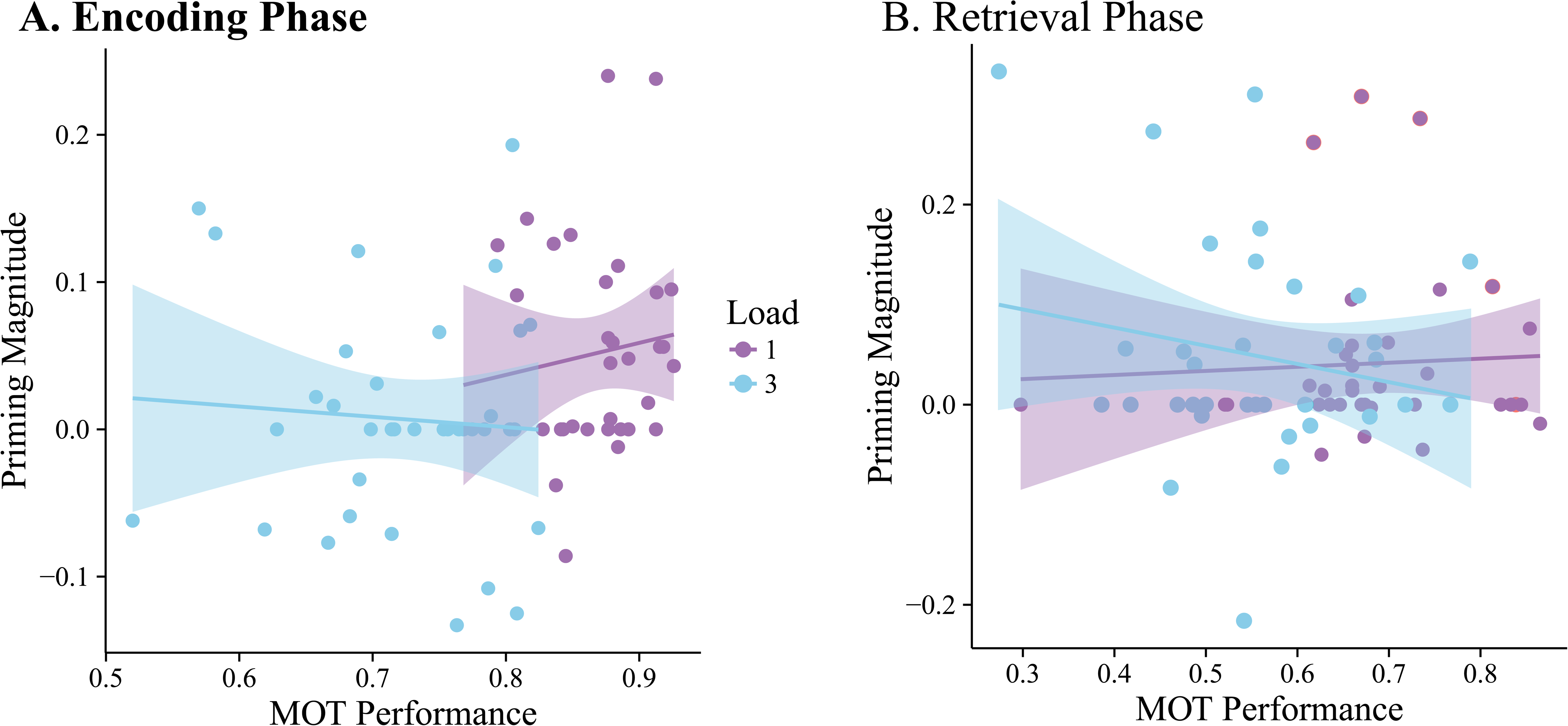
Predictability of priming magnitude based on single task MOT performance. The lack of a correlation between priming magnitude and MOT task performance in either the Encoding or Retrieval phases suggests that being good at one task does not predict performance in another task.

## Discussion

We utilized a dual-task experiment to determine whether syntactic processing required language-specific or domain-general resources. To measure syntactic processing, we used a syntactic priming paradigm. We modulated the amount of attentional resources available by using a motion object tracking (MOT) task, which is commonly used in the literature in this context (Allen et al., 2004; Fougnie & Marois, 2006; Postle et al., 2005). In addition to modulating attention, we manipulated whether the MOT task was performed concurrently to the encoding of syntactic information (Encoding phase), or concurrently to the retrieval of previously processed syntactic information (Retrieval phase). A comparison between the Encoding phase and the Retrieval phase experiment will determine whether these two phases use similar attentional resources.

Accuracy in the MOT task was significantly reduced in the dual-task condition compared to the single task condition. This is consistent with the claim that the MOT and the Encoding/Retrieval phases of the syntactic priming task tap into the same resources (Kinchla, 1992).

Interestingly, a drop in syntactic priming magnitude was not seen for conditions in which the participants had to track one or three balls compared to conditions in which they had to track no balls (i.e., single-versus dual-task conditions). As the MOT task was always presented first, this suggests that the language task was given the priority when it came to resource allocation (Lavie & Tsal, 1994), even though we never told participants to focus more on the language task compared to the MOT task. This result is at odds with another dual-task study that looked at language production and comprehension while driving and found that driving was given the priority over language (Bock, Dell, Garnsey, Kramer, & Kubose, 2007; Kubose et al., 2006). However, the authors explained this as driving being given the priority due to the life-threatening nature if it wasn’t. This, together with our results, suggests that the natural preference of one task over another is highly sensitive to context.

The drop in performance for the MOT task was significantly lower for the Retrieval phase compared to the Encoding phase, suggesting that either speaking requires more resources than listening, or speaking has more in common with visuospatial attention than listening. We did, however, see a robust *increase* in priming magnitude in the Encoding phase for Load (1). This increase was more than double the priming magnitude observed in the control phase (4.6% versus 10.4%).

This enhancement is not a result we predicted to find. Although post-hoc and speculative, we find that our result is consistent with a phenomenon known in the field of attention research as the attentional boost (Swallow & Jiang, 2011, 2014). When a target appears, no matter if it is a frequent target or not (Makovski, Swallow, & Jiang, 2011; Swallow & Jiang, 2012) attention to this target leads to widespread increases in perceptual processing. The attentional boost hence suggests that there are resources left over in reserves that are allocated when the target appears (dual-task interaction model; Swallow & Jiang, 2013). This is consistent with the results we observe in our current study: Participants have assigned the language task as the goal-relevant task and have assigned the majority of their resources to it. However, the appearance of a ball to track causes the participants to increase their perceptual processing while they are trying to keep track of this one ball. This causes them to encode the picture and the sound file better compared to conditions in which they have no balls to track. When the participants have three balls to track, however, the MOT task already demands extra resources as now the participants have to encode three balls, not one. This means that there are no extra resources to recruit for the boost they would have otherwise received. This enhancement does not occur for the retrieval phase, because they are retrieving stored information, not perceiving anything when they are conducting the MOT task. This does not rule out that the participants in the Retrieval phase do not have a boost during the MOT task, but there is nothing to perceive except the picture as the participants in the Retrieval phase only conduct the MOT task when they are describing the picture. There is also no effect on the active structures, as syntactic priming only occurs for the infrequent structures (inverse preference effect; Bock, 1986; Bock, Loebell, & Morey, 1992; Ferreira & Bock, 2006). This explanation provides an interesting basis for further research into language and the attentional boost.

Moreover, this increase in priming magnitude was not only seen in the priming magnitude of the Load (1) condition in the Encoding phase, but also in the catch rate for the same condition, providing converging evidence. This suggests that the increase in priming magnitude is not only limited to enhanced memory for the syntactic structure, but could also be an enhancement in the integration of syntactic structure and visual information. The effect of attention on integration has been observed before in the McGurk effect (McGurk & MacDonald, 1976). The illusion is driven by the integration of audio and visual streams, yet under high attentional load this illusion breaks down, as the integration is not possible with such limited attentional resources (Alsius, Navarra, Campbell, & Soto-Faraco, 2005).

Our results are also interesting in relation to a more general application: Multi-tasking while driving as driving also involves constant spatial attention similar to the MOT task. Previous studies on language and driving have shown that it is not the handling of a cell phone that is dangerous while driving, it is the act of conversing itself (see Strayer, Watson, & Drews, 2011 for a review). Research on memory for language has shown that recall accuracy for a recently comprehended short story is significantly impaired if done while driving (Bock et al., 2007; Kubose et al., 2006). However, if the participant recalls the story while not driving (although it was still told when the participant was driving), there is no significant difference compared to when the participant heard the story while not driving. This result is inconsistent with our study as it suggests that it is not the comprehension/encoding of information that is affected, it is the retrieval of that information. However, many aspects of the Kubose and colleagues study have been explained as driving being a highly practiced and therefore semi-automatic process. Therefore, perhaps our results are a better reflection of beginner drivers where the task of driving is not as highly practiced.

Overall our results show that language gets priority in terms of assignment of the available resources when it is shared with a non-language (perceptual) task. The decrease in performance we expected to see if syntactic processing and the MOT task tap into the same resources was only seen in the performance of the MOT task, not in priming magnitude. Nevertheless, it does suggest that syntax and MOT tap into the same resources, in this case domain-general resources as a decrease in performance was seen.

Although the MOT task was selected due to its “pure” attentional manipulation, it is still a perceptual task, and hence does tap into domain-specific resources, such as visuospatial attention, in addition to domain-general resources. Therefore, our results could also be interpreted as evidence that language taps into visuospatial attention, and hence the interference with the MOT task. There have been suggestions that when people read or listen, they create spatial references in their mind (Fincher-Kiefer, 2001; Fincher-Kiefer & D’Agostino, 2004). However, as we give the participants the picture of the description, we are hard-pressed to suggest that our participants additionally created unnecessary spatial references which could have interfered with the MOT task. Hence we do not believe that the task interference was caused by shared visuospatial resources, but rather shared domain-general resources.

In summary, our results suggest that syntactic processing is not an automatic process, and does require attention to operate. Even though we do not see a drop in priming magnitude in the language task, we do see a drop in the MOT task performance, meaning that the MOT task had less resources to complete the task accurately. This could only have occurred if another task was tapping into the same pool of resources. It also suggests that language receives priority in terms of assignment of the available resources when it is shared with a non-language (perceptual) task. The attentional boost effect seen in the Load (1) Encoding phase condition is interesting and has not been observed before for modality independent processes. It poses the question if this effect can be seen for other non-automatic language processes and what role this effect could play in language comprehension.

## Acknowledgements

We thank Peter Hagoort, Yingying Tan, and Kelly Garner for their valuable insights on previous versions of this manuscript.

